# Divergent east-west lineages in an Australian fruit fly associated with the Carpentaria basin divide

**DOI:** 10.1101/2022.10.04.510875

**Authors:** Chapa G. Manawaduge, Anthony R. Clarke, David A. Hurwood

## Abstract

*Bactrocera jarvisi* is an endemic Australian fruit fly species (Diptera: Tephritidae). It occurs commonly across tropical and subtropical coastal Australia, from far-northern Western Australia, across the ‘Top End’ of the Northern Territory, and then down the Queensland east coast. Across this range, its distribution crosses several well documented biogeographic barriers. In order to better understand factors leading to the divergence of Australian fruit fly lineages, we carried out a population genetic study of *B. jarvisi* from across its range using genome-wide SNP analysis, utilising adult specimens gained from trapping and fruit rearing. Populations from the Northern Territory (NT) and Western Australia were genetically similar to each other, but divergent from the genetically uniform east-coast (=Queensland, QLD) population. Phylogenetic analysis demonstrated that the NT population derived from the QLD population. We infer a role for the Carpentaria Basin as a biogeographic barrier restricting east-west gene flow. The QLD populations were largely panmictic and recognised east-coast biogeographic barriers play no part in north-south population structuring. While the NT and QLD populations were genetically distinct, there was evidence for the historically recent translocation of flies from each region to the other. Flies reared from different host fruits collected in the same location showed no genetic divergence. While a role for the Carpentaria Basin as a barrier to gene flow for Australian fruit flies agrees with existing work on the related *B. tryoni*, the reason(s) for population panmixia for *B. jarvisi* (and *B. tryoni*) over the entire Queensland east coast, a linear north-south distance of >2000km, remains unknown.

## Introduction

In the western Pacific, the Dacini fruit flies (Diptera: Tephritidae) are a super-diverse clade of more than 400 species, predominantly occurring within the genus *Bactrocera* Macquart (1, 2). Larval breeding occurs in fleshy fruits and most species are restricted to tropical rainforests, the presumed ancestral habitat of the genus (1). A few species, such as *Bactrocera tryoni* (Froggatt) and *B. frauenfeldi* (Schiner) are damaging pests of horticulture (3, 4), but most species are non-pestiferous.

The evolutionary drivers for the extensive speciation of *Bactrocera* are not obvious. As fruit specialists, diversification could follow ecological specialisation onto new hosts (5). However, *Bactrocera* is unusual in the high levels of polyphagy exhibited by many species (6) and regional radiation has not been associated with increasing levels of host-use specialism (7). The case for allopatric speciation is also difficult to develop, as *Bactrocera* communities are typified by high alpha-diversity and low beta-diversity, that is a single location has many co-occurring species but there is little species turn-over between locations (8, 9). Starkie et al (7) found evidence of species radiations associated with the movement of flies from New Guinea into South Pacific islands and Australia which is supportive of allopatrically driven divergence; but within Australia there was no evidence for the east-coast biogeographic barriers (10) being associated with clade divergence, which is not supportive of allopatric speciation. More research is thus clearly needed to better understand the drivers of divergence in the Dacini.

Population genetics seeks to understand population structuring and divergence within a species and is thus another tool for untangling the drivers of initial divergence within *Bactrocera* species. Population structuring in tephritid species is generally weak (11, 12), but nevertheless population-level divergence within tephritid species has been linked with differential host use (13, 14), increasing geographic distance (15, 16), and gene flow restriction associated with biogeographic barriers (17, 18). Within Australia, population genetic studies have only been applied to the pest *B. tryoni*, with the general pattern of findings for this species being little or no genetic structuring across the vast distribution (>2000km) of the endemic east coast population, but with structuring from east to west across Australia’s tropical ‘Top End’ (19–22). However, interpreting these results is difficult as some samples from the Northern Territory (NT), and exclusively from Western Australia (WA), include a taxonomically different species, *B. aquilonis* (May). Whether *B. aquilionis* represents a sister species to *B. tryoni*, or a still diverging lineage of *B. tryoni* is unclear from molecular data (22), but either scenario demonstrates the potential for allopatric divergence to drive the diversification of Australian fruit flies. However, with only one species studied, how general a pattern of east-west divergence might be for Australian fruit flies is unknown.

*Bactrocera jarvisi* (Tryon), Jarvis’ fruit fly, is an Australian Dacini species endemic to northern and eastern coastal regions of Australia (23). It is a polyphagous species, having been reared from 83 wild and commercial hosts (24). While a recognised pest of the Northern Territory mango industry (25, 26), its major native host over most of its range is considered to be the small woodland tree, *Planchonia careya* (F.Muell.) R.Kunuth (Lecythidaceae), commonly known as cocky apple (27, 28).

Our primary aim here was to explore whether biogeographic barriers may be associated with population structure in *B. jarvisi*. In this study, we investigated the genome-wide single nucleotide polymorphism (SNP) variation found in *B. jarvisi* adult samples from across most of the species’ geographic distribution to identify where population differentiation may exist. Additionally, to determine if there is any evidence for host-associated divergence, we analysed SNP variation from individuals reared from commercial and native host fruit species.

## Materials and methods

### Sampling

Individuals of *B. jarvisi* were collected between December 2020 and February 2022, either using zingerone lure traps in the field or rearing from infested fruits in the laboratory (S1 Table). The geographic range coverage included samples from the very west of the known *B. jarvisi* distribution in Western Australia (WA), then across the Northern Territory (NT) to the east coast of Queensland (QLD); while latitudinal sampling was undertaken from Cooktown in far-north Queensland (FNQ) to its southern most distribution in far northern New South Wales (NSW) (Fig 1). From zingerone trapping, 188 individuals were sampled from 18 locations across the species’ geographic distribution. Further, 81 individuals were reared from five hosts (native hosts: cocky apple, white bush apple (*Syzygium forte*); introduced hosts: mango (*Mangifera indica*), plum (*Prunus domestica*), guava (*Psidium guajava)*). All samples were preserved in 100% ethanol and stored at −20°C until processing.

**Fig 1.**
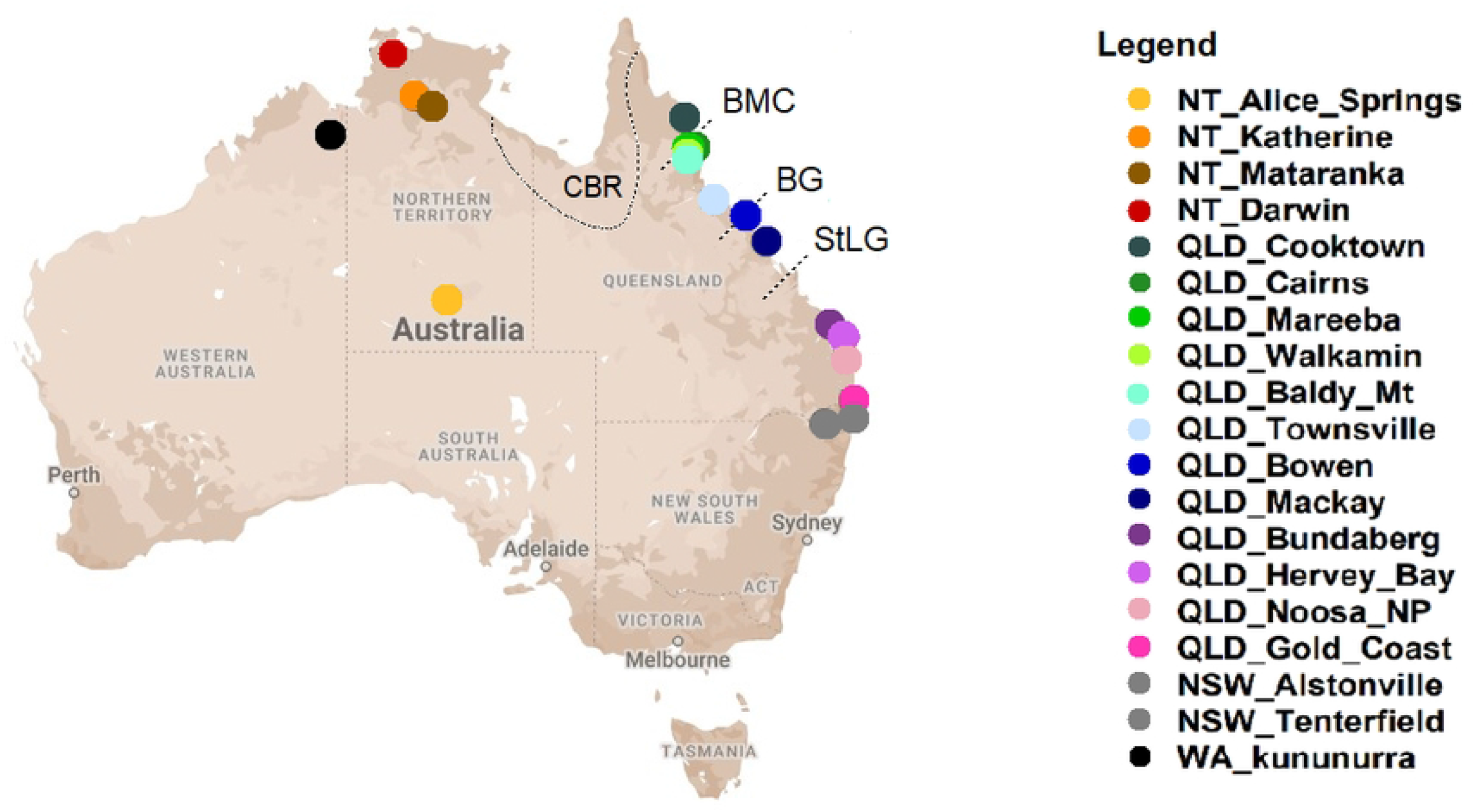
Map illustrating the sampling locations of *Bactrocera jarvisi*. Each colour represents a different sampling location. Dashed lines indicate the locations of recognised biogeographic barriers, Carpentaria Basin Region (CBR), The Black Mountain Corridor (BMC), the Burdekin Gap (BG), and the St Lawrence Gap (StLG). Marking for CBR is generalised from several sources that have attempted to characterise this barrier based on the distributions of a variety of taxa).

### DNA extraction and genotyping

For DNA extractions, either the head and legs or the full body without the abdomen was used depending on the samples’ freshness and quality. DNA was extracted using QIAGEN DNeasy® Blood & Tissue Kit following the manufacture’s protocol with only slight modifications. DNA samples were screened for quality and quantity, using both gel electrophoresis and Qubit assay, and then sent to Diversity Arrays Technology Pty Ltd, Canberra (DArT P/L), for DArTseq high-density genotyping. The restriction enzyme combination, PstI/SphI, was used by DArT P/L for the current *B. jarvisi* dataset, and the fragments (up to 90bp) were sequenced on an Illumina Hiseq2500 as single end reads.

### SNP analysis

Sequences were processed in proprietary DArT analytical pipelines, and the results of the SNP calling were returned as a matrix. These SNPs results were viewed and assessed for quality and informativeness using the DArTR package (29) in the R platform (30). Low quality loci (< 80% call rate and < 95% reproducibility) and individuals (< 80% call rate) were filtered out. Monomorphic loci were also removed before further analysis.

### Genetic relationship and admixture analyses

A principal component analysis (PCA) was performed on the filtered data set using DArTR package, to visualise the overall genetic relationships of the individuals among locations. Because the data set consisted of some individuals that were reared from infested fruits, pairwise kinship coefficients were estimated using the maximum likelihood estimation method in the R package SNPRelate (31). Preliminary analysis of these data has suggested that there may be some analytical artefacts driven by the inclusion of individuals that may be very closely related to each other because they have been reared together. Therefore, only the trapped samples were used to investigate the genetic structure and relationship across the distribution. Fruit-reared samples were analysed separately.

### Admixture analysis

To estimate the individual admixture coefficients, the R package, LEA (32), that implements a sparse, non-negative matrix factorisation algorithm (sNMF) to estimate ancestry coefficients from large genotypic matrices and to evaluate the number of ancestral populations (K), was used. The entire data set was run for K = 1–20, with 100 repetitions per each K value. The value of K that corresponded with the lowest cross-entropy criterion was selected as the value that best explained the results (32).

### Phylogenetic analysis

For the phylogenetics analysis, the filtered SNP dataset was concatenated and stored in a fasta format file, in which SNPs were converted into base pairs following IUPAC ambiguity codes for heterogeneous SNPs using the DArTR package. Phylogenetic trees were then constructed using a maximum likelihood method in IQ-TREE 2.12 (33). The model finder option coupled with the ascertainment bias correction was used to select the best fit model, and the phylogenetic trees were reconstructed with 10,000 ultrafast-bootstraps and 1,000 bootstrap replicates for the SH-like approximate likelihood ratio test. An additional step to further optimise UFBoot trees by nearest neighbour interchange based directly on bootstrap alignments was also incorporated to reduce the risk of overestimating branch supports due to severe model violations. Finally, the support of alternative topologies was tested using the RELL approximation method (34) in IQ-TREE. All the tree files were viewed and edited in Figtree v.1.4.3 (http://tree.bio.ed.ac.uk/software/figtree/).

## Results

A total of 29,286 binary SNPs was obtained for 269 individuals with 36% of missing data. Loci call rate ranged from 20–100% with an average of 64%, and scoring reproducibility ranged from 93– 100% with an average of 99.7%. After filtering out low quality data, the final dataset retained 9,769 SNPs for 260 individuals with only 5% missing data. The average loci call rate and reproducibility were 94% and 99% respectively.

The initial principal component analysis of the combined data set (i.e., adult trapping + fruit rearing) revealed some level of genetic structure within *B. jarvisi* (Fig 2). Along the first two PCA axes, that account for 8.2% of the sample variance, the majority of individuals from the NT form a tight cluster and were largely divergent from individuals from the east that formed a more diffuse group. However, a few individuals sampled from NT clustered with individuals from the east, and vice versa. For the individuals obtained from fruit rearing, no clear genetic pattern was found with respect to host, but there was a degree of family structuring (Fig 3) with close genetic relatedness among the flies reared from a single host. This was supported by pairwise kinship coefficient analysis (S1 Fig). Close family relatedness in host-reared material is not unexpected, as individuals may be derived from an individual egg clutch (i.e., they have the same mother).

**Fig 2.**
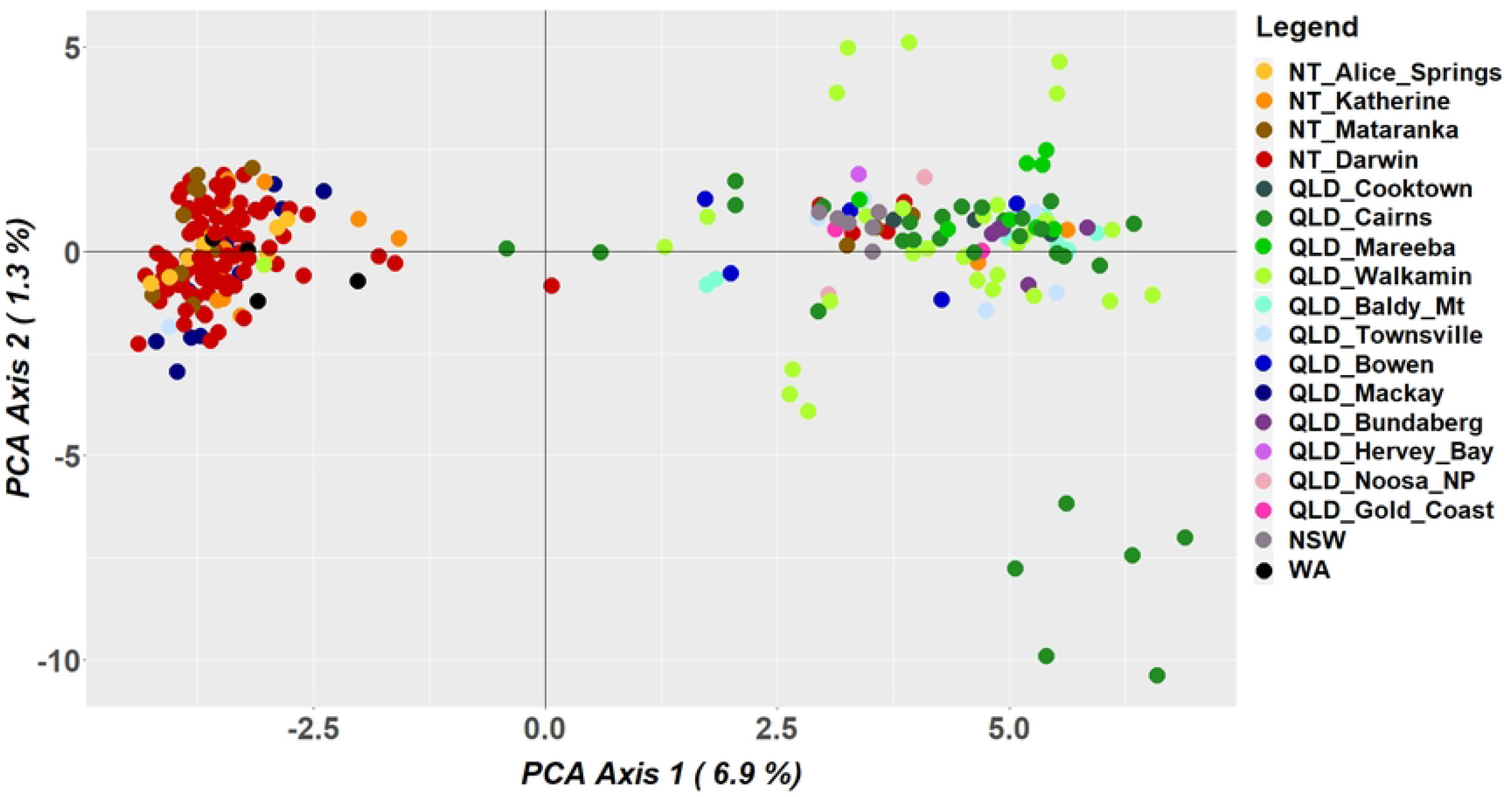
Scatter plot from a principal component analysis of *Bactrocera jarvisi* genome-wide SNP data containing all individuals sampled from trapping and host-rearing. Each colour represents a different sampling location corresponding with Fig 1.

**Fig 3.**
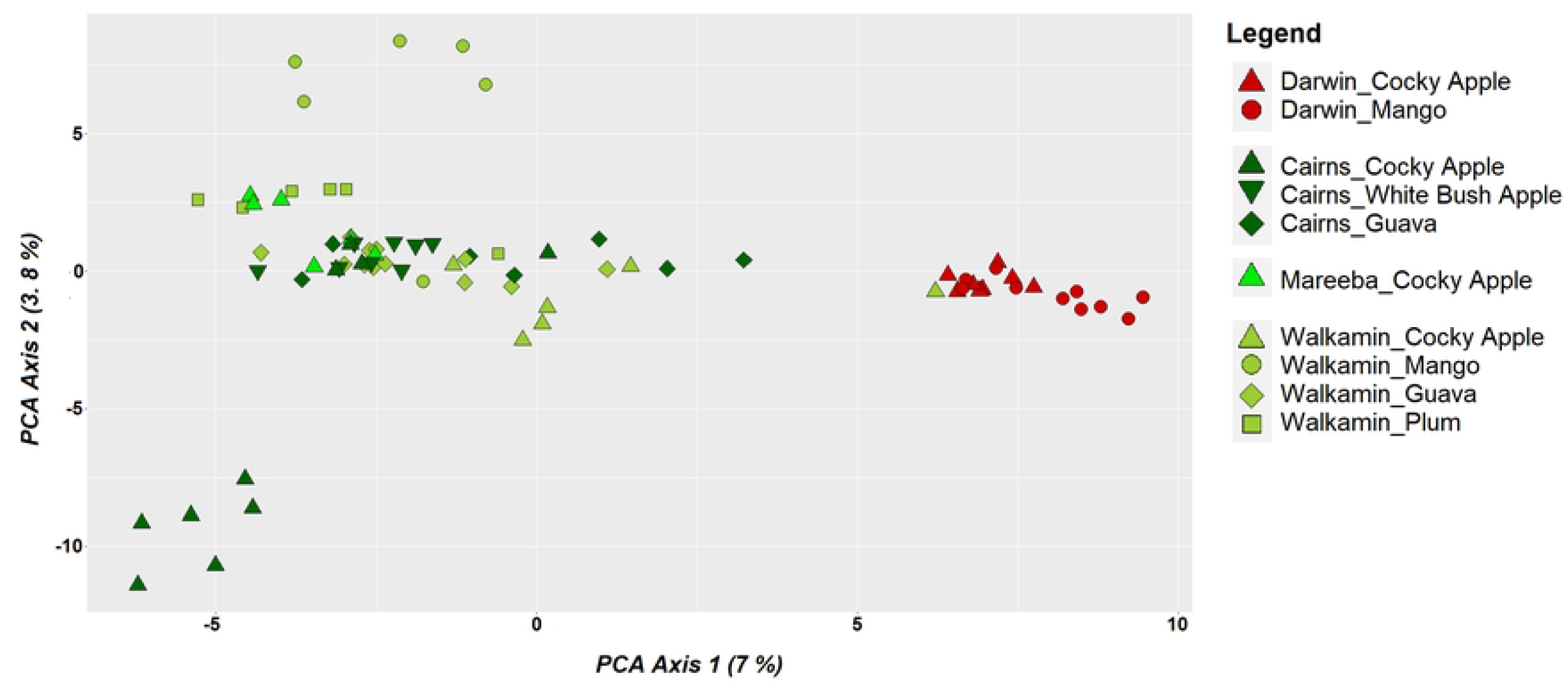
Scatter plot from a principal component analysis of *Bactrocera jarvisi* genome-wide SNP data containing only individuals reared from host fruits. Each colour represents a different sampling location corresponding with Fig 1 and each shape represent a different host species.

After removing the host-reared individuals and filtering out low-quality data, the data set containing only trapped samples consisted of 182 individuals with 8,102 SNP loci with a 94% average loci call rate and 5% missing data. The admixture analysis of these data supported K = 3 (i.e., 3 major groups) as the values corresponding with the lowest cross-entropy from the sNMF algorithm in LEA (Fig 4) and supports the existence of two major groups along with one additional small cluster. Almost all the samples from the NT and WA, along with a few FNQ samples belong to a separate genetic group, hereafter referred as ‘*NT cluster*’, whereas the majority of the east coast samples (QLD and NSW), together with a few individuals from Darwin, represent a second genetic group, here after referred as ‘*QLD cluster*’. A third group contained only six individuals that came from Darwin. However, when viewing the phylogenetic tree (Fig 5) these individuals do not cluster together as may be expected. Therefore, we assumed that this third cluster is an artefact of the algorithm used to search for the optimum number of genetically differentiated groups. As such we consider them to be members of the NT cluster.

**Fig 4.**
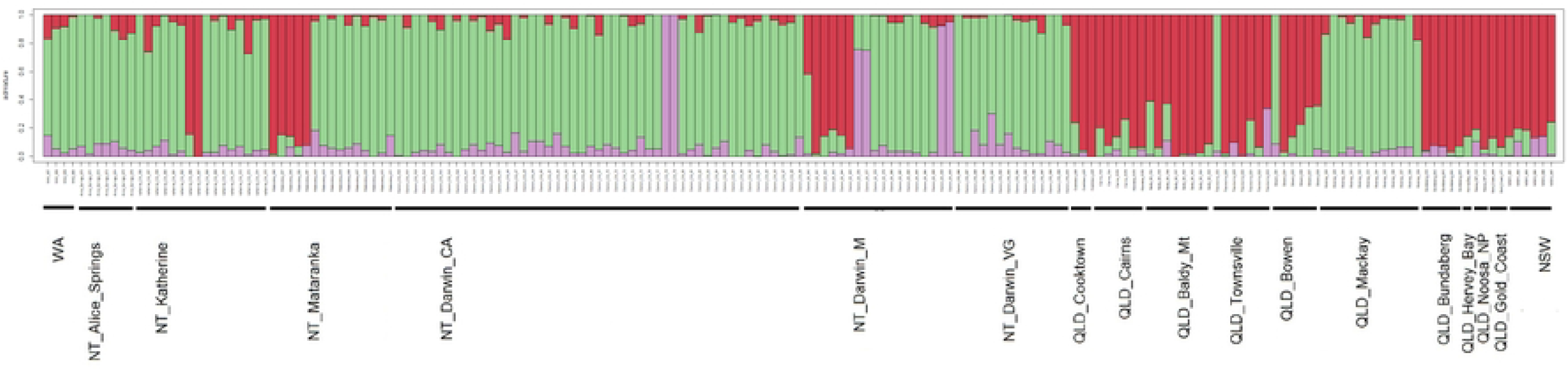
Estimates of *Bactrocera jarvisi* admixture proportions inferred with sNMF using genome-wide SNP data. Each bar is an individual and displays the relative assignment probability to each of three identified genetic groups.

**Fig 5.**
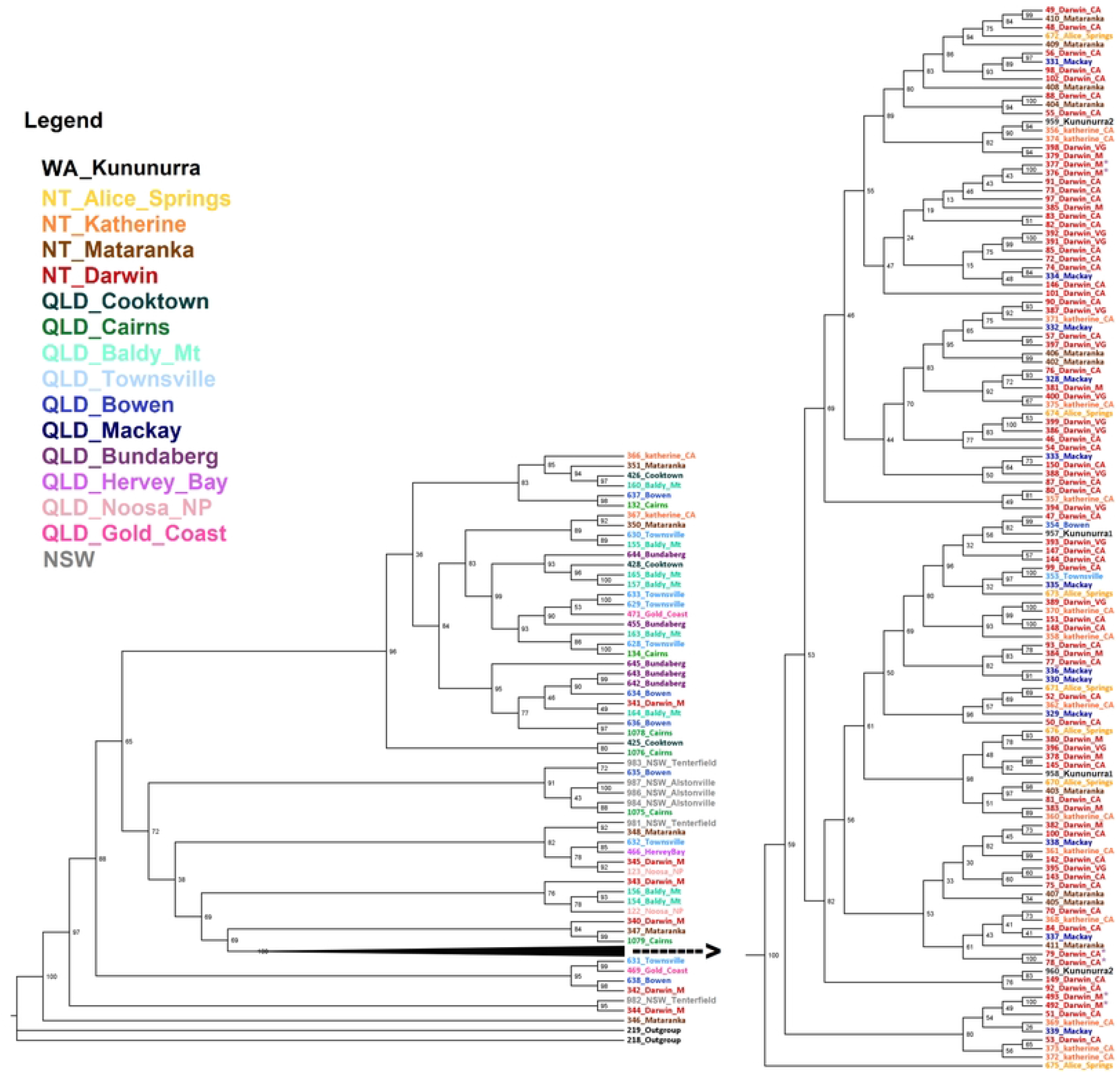
Phylogenetic tree based on genome-wide SNP data for populations of *Bactrocera jarvisi* collected from across the species’ range. [SNP maximum likelihood (IQ-TREE) consensus tree from 10,000 ultrafast-bootstrap replicates. Each colour of the tip labels (sample IDs) corresponds with the location colours given in Fig 1, and the numbers at each node represent the ultrafast-bootstrap support. *Six individuals that represent the third group in the admixture analysis (Fig 4)]

Phylogenetic analysis in IQ-TREE required exclusion of constant genetic sites, that resulted in a final dataset with 6,519 SNPs, of which 3,767 were parsimony-informative. For the maximum likelihood analysis, the SYM+ASC+R10 model was selected as the best fit model according to the Bayesian information criterion of 30 candidate DNA models, analysed by the model-finder option in IQ-TREE. *Bactrocera cacuminata* was considered as the out group, and the resulting phylogenetic tree highly supported *B. jarvisi* (100% ultrafast-bootstrap support) as a monophyletic group (Fig 5, S2 Fig). However, the individuals from the *NT cluster* are recently evolved from a single linage. Individuals of this clade form a relatively tight cluster opposed to the large and defuse *QLD cluster*, indicating relatively low genetic variation compared to the individuals from the *QLD cluster* that correspond with the older lineages.

## Discussion

The results here represent the first comprehensive study on the genetic structure of *B. jarvisi* across its distribution. After the *B. tryoni/B. aquilonis* species complex (22, 35), this is only the second Australian fruit fly species for which population genetics has been completed. The major findings were that *B. jarvisi* exhibits: i) significant genetic structure between QLD and NT populations, ii) a lack of differentiation across the east coast (QLD+NSW), as well as across northern Australia (NT+WA), iii) some evidence for translocation in both directions and iv) no evidence of host-associated divergence. Most population genetic studies undertaken on Dacini fruit flies have found only weak genetic structuring over very large geographic distances (36–39); however, in some cases, structure has been detected that can be associated with known biogeographic barriers (17, 18). In the current study we have found both; some structuring which can be associated with a biogeographic barrier, but also a lack of structure despite biogeographic barriers and very large distances between sampling sites. The following discussion addresses these issues further.

### Role of biogeographic barriers in *B. jarvisi* population structuring

A number of biogeographic barriers have been identified along the northern and eastern coastlines of Australia (Fig 1) (10, 40) and these barriers have been shown to influence the dispersal of many taxa including amphibians (41), reptiles (42), mammals (43, 44), birds (45), insects (46) and plants (47). To date however, there is no evidence to suggest that the east-coast biogeographic barriers have had any significant influence on the historical movement of Dacini (7, 10), and this is again the case for *B. jarvisi*. However, we found a significant differentiation of *B. jarvisi* populations between QLD and NT. Given the homogeneous nature of east coast *B. jarvisi* populations over >2000kms, the observed level of differentiation between QLD and NT populations cannot be accounted for by distance alone. If this is the case, we need to look for a barrier to dispersal, whether physical or climatic, between NT and QLD.

The Jurassic-Cretaceous intracratonic Carpentaria Basin is a well-documented phylogeographic barrier in northern Australia that extends from QLD to the NT (Fig 1) (48). This barrier has played a major role in differentiating many taxa including the avian fauna of northern Australia (49). In their study of *B. tryoni*, Popa-Baez et al (22), also found significant genetic structure between *B. tryoni* and *B. aquilionis* along the east-west transect of northern Australia. Whether *B. aquilionis* represents a sister species to *B. tryoni* or a still-diverging lineage of *B. tryoni* is debatable, but nevertheless these lineages clearly exhibit restricted gene flow (19, 22). The Carpentaria Basin is a semi-arid region extremely poor in vegetation (49), and currently, there are no records of either of these two species, or indeed any Dacini fruit fly species in this region (24, 50). Together, with the prior work on the *B. tryoni/B. aquilonis* system, our study strongly suggests that the Carpentaria Basin acts as a biogeographic barrier to Dacini and plays a role in the allopatric diversification of fruit fly species in Australia.

### Role of human mediated dispersal in *B. jarvisi* population structuring

While the influence of the Carpentaria barrier in constraining dispersal of *B. jarvisi* populations is evident, we still detected instances of high gene flow between NT and QLD in the data. Specifically, the Mackay sample is made up of ‘NT’ genotypes and a few individuals from Darwin and Mataranka in the NT have a QLD genotype. The most parsimonious explanation for this anomaly is human-mediated translocation. It is interesting to note that the whole Mackay sample was made up of introduced NT types with little evidence for introgression with local QLD types. While our study did not have the scope to address such issues, questions arising regarding non-random mating among lineages and competitive exclusion between a local and introduced lineage should be investigated.

There was a noted lack of genetic differentiation among *B. jarvisi* samples along the east coast (QLD/NSW), a pattern also seen in *B. tryoni* (22), with the exception of the Mackay population. This may be evidence for repeated, human-assisted movement of fruit flies to that region. Human carriage of fruit fly infested fruit with resultant panmixia has been used as an explanation for the lack of genetic structuring in numerous pest fruit fly species (38, 51–53). While often proposed, this explanation must assume for *B. jarvisi* and *B. tryoni*, that human movement of infested fruit is sufficiently extensive to account for the lack of genetic differentiation across >2000km and to hide the historical genetic signatures of multiple, well documented biogeographic barriers. While we consider this unlikely in the well-managed Australian horticultural system, particularly for *B. jarvisi* which is a lesser pest, the hypothesis is unfortunately very difficult to test. Fortunately, this is not the case for the corollary of the hypothesis. If human-mediated movement does explain the lack of genetic structure for pest species such as *B. tryoni* and *B. jarvisi*, then it is logical that a non-pest species would not be exposed to human-mediated dispersal and so should show a genetic structure more consistent with historical biogeography, current landscapes, and host-species’ distributions. To better understand the drivers of Dacini population genetic structuring, we consider population genetic studies on wide-spread, non-pest species to be a priority.

### Role of host in *B. jarvisi* population structuring

Population level divergence of some tephritid fruit flies through adaptation to novel hosts has been documented (13, 14, 54). For example, genetic differentiation of *Rhagoletis pomonella* between co-occurring hosts, hawthorn (the native host) and domestic apples, has been reported with a reduction in gene flow and sympatric divergence (13, 14, 54). However, the genetic differentiation observed across the different host species in *B. jarvisi*, while consistent with family structuring, provided no evidence for sympatric, host-associated divergence in the species.

The close association between *B. jarvisi* and its native host, cocky apple, has long been known (27, 55, 56) and it has been speculated that cocky apple abundance plays an important role in shaping and maintaining *B. jarvisi* populations, even in the presence of alternate commercial hosts such as mango (26, 27, 57). If breeding populations of *B. jarvisi* were dependent on cocky apple for the primary maintenance of populations, it could be argued that samples collected from cocky apple should be of the same genetic type as those samples collected from nearby but different host(s). However, such similarity or relatedness was not found among the flies from Walkamin Research Centre (FNQ) that were reared from four different host fruits: cocky apple, mango, guava and plum. Kinship analysis revealed no close relatedness between the flies reared from cocky apple and other hosts in sympatry.

## Conclusion

This is the first study of the genetic structure of *B. jarvisi*. We determined that: *B. jarvisi* exhibits significant genetic structure between QLD and NT populations, although there is evidence for recent translocation in both directions; a lack of genetic differentiation among the east-coast populations; and no evidence of host association divergence. Our study suggests that while the biogeographic barrier of the Carpentaria basin may be playing a role in the diversification of *B. jarvisi* in northern Australia, there is no evidence for the biogeographic barriers along the east coast of QLD influencing the genetic structure of *B. jarvisi*. Whether human-mediated dispersal could account for the lack of genetic differentiation across >2000k and several well-established biogeographic barriers is questionable. To test this a corollary of the human-mediated dispersal hypothesis, we should target a population genetic study on a non-pest species That is less likely to be moved by humans. Within Australia, the wild tobacco fly, *B. cacuminata* (Hering), is an abundant, widely distributed non-pest species suitable for such a study.

## Acknowledgments

We thank for the Commonwealth Government for its support and acknowledge the overall project leadership of Mr Peter Leach (Department of Agriculture and Fisheries, Queensland). Dr Thilini Ekanayake (Department of Primary Industry and Resources, Northern Territory), Dr Solomon Balagawi (New South Wales Department of Primary Industries), Dr Melissa Starkie and Mr Stefano De Faveri (Department of Agriculture and Fisheries, Queensland), Dr Touhidur Rahman (Department of Primary Industries and Regional Development, Western Australia), all supplied material without which the project would have been impossible to complete. We thank them and their institutions for their support and engagement.

## Supporting information

S1 Table. Details of the samples used in the study. (Sampling method= T: Trapping, R: Rearing)

S1 Fig. Heat map showing the pairwise kinship coefficients between the reared individuals. (Kinship coefficient: 0.5 = monozygotic twins, 0.25=full siblings, 0.125=half siblings, 0.0=unrelated)

S2 Fig. SNP maximum likelihood (IQ-TREE) consensus tree from 10,000 ultrafast-bootstrap replicates showing the true branch lengths. Each colour of the tip labels (sample IDs) corresponds with the location colours given in Fig. 1. The numbers at each node represent the ultrafast-bootstrap support.

## Notes

### Competing Interest Statement

The authors have declared no competing interest.

